# Comprehensive Characterization of tRNA by Ultra High-Performance Liquid Chromatography High-Resolution Accurate Mass Spectrometry

**DOI:** 10.1101/2023.06.07.544034

**Authors:** Robert L Ross, Ryan Cowley, Keeley Murphy, Roberto Gamez, Paul R Gazis, Jennifer Sutton, Min Du

## Abstract

Transfer ribonucleic acid (tRNA) are the smallest RNA in the translational triune and contain the greatest density of post-transcriptional modifications than any other RNA types in the cell. Due to the size of tRNA studying these modifications usually entails enzymatic digestion followed by liquid chromatography tandem mass spectrometry (LC-MS/MS). Here we report an advancement in Intact Mass Analysis for identification of tRNA through deconvolution of high resolution accurate mass spectrometry facilitated using a secondary alkylamine as an ion pairing reagent during reverse phase chromatography. We identify in isolated and total *S. cerevisiae* tRNA 3’ CCA variations and show that most tRNA lack a 3’ adenosine with the lesser abundant species having the expected CCA termini. We identify a previously unknown stable demethylated Wybutosine intermediate for tRNA^PHE^ and identify low abundant contaminating tRNAs in an isolated tRNA^PHE^ sample. Confirmation of identities was verified through traditional mass spectrometric nucleoside and mass mapping experiments.

## INTRODUCTION

Post transcriptional modifications of RNA range from simple methylations to more intricate chemistries resulting from extensive enzymatic pathways.^1–3^ Over 140 chemical modifications to the canonical base and/or sugar of RNA have been reported.^4^ These modifications act as structural elements for the RNA,^5–7^ assist with translation,^8,9^ and act as determinates for RNA-Protein conjugation^10^ that has been shown to regulate gene expression.^11–13^ By far the greatest density of modification is found in the tRNA, with an average of 13 modifications per tRNA.^14,15^ Numerous disease states such as cancer,^16–18^ neurological disorders,^19–21^ birth defects^22,23^ and cardiovascular disease^24–26^ have been linked to mutations within the genes for modification enzymes highlighting their importance in biological processes.

Analytical methods used to study these modifications began with anion exchange chromatography.^27–29^ As technology improved chromatography was hyphenated to mass spectrometry^30,31^ affording the ability for structural characterization. Techniques from Sanger Sequencing were adopted as a strategy for mapping modification to their sequence loci.^32^ Further advances yielded RNA-Seq^33,34^ where through reverse transcription the RNA is converted back to complementary DNA (cDNA) and then sequenced. This approach suffers from the loss of modification status as the polymerase enzyme cannot read through the modifications generating hard stops in the cDNA. When sequenced these hard stops give location of modification, but not identity. Chemical derivatization of the RNA followed by sequencing is shown to identify specific modifications such as methylations^35^ or pseudouridinilation,^36^ and while powerful, these methods are modification specific and cannot identify all modifications present on the strand in a single assay.

The gold standard in analytical technology for RNA identification and characterization remains steadfast in the realm of liquid chromatography online with tandem mass spectrometry (LC-MS/MS).^37–39^ LC-MS/MS is a direct measurement of the RNA, as such it provides the best method for characterization of post transcriptional modifications. Historically, due to the size of the biomolecule, tRNAs have been subjected first to enzymatic digestion to produce smaller oligonucleotides^32,40,41^ which are then separated chromatographically before being analyzed by the mass spectrometer. When the oligonucleotide is subjected to collision induced dissociation (CID), the energy from the collision is distributed over the length of the biopolymer fragmenting the oligonucleotide at the weakest bond, usually at the diester bond of the terminal phosphate.^42^ This fragmentation happens sequentially, where a single nucleotide is lost at each end of the biopolymer for each CID event. The resulting fragmentation spectra allows the sequence of the oligonucleotide to be observed at nucleotide level where each nucleotide in the sequence can be viewed sequentially in the spectra.

One of the challenges with LC-MS/MS of oligonucleotides is the hydrophilicity of the biopolymer. The electrospray process transitions the RNA from the liquid phase into the gas phase as an ion which can be detected by the mass spectrometer. The use of ion pairing reagents are used to mask the electronegative backbone for chromatographic retention and to assist rapid desolvation of the oligonucleotide in the electrospray plume.^43^ After transitioning from the droplet to the gas phase the oligonucleotide exists as a series of multiply charged ions with associated mass to charge ratios (*m/z*) and results in a complex spectrum. Through a process called deconvolution a mathematical algorithm reconstructs the acquired data into interpretable results. Deconvolution works best when the oligonucleotide presents as isotopically resolved peaks in the spectrum, therefore the transition from the liquid phase to the gas phase is critical for clean isotopically resolved spectra. When the oligonucleotide does not fully desorb from the electrosprayed droplet as a naked ion, the droplet will evaporate resulting in metal adduction^44^ to the oligonucleotide which masks the signal.^45^ Poor desorption from the droplet has resulted in the inability to extensively study longer oligonucleotides and their modifications by mass spectrometry.

Here, using Ultra High Performance Liquid Chromatography High Resolution Accurate Mass Spectrometry (UHPLC-HRAM-MS) we show the ability to identify tRNA through Intact Mass Analysis (IMA) facilitated using a hydrogen bond donor alkylamine. Through IMA we verify past identity of tRNA^PHE^ having a CpCp 3’ end and show that most tRNA end with this truncation. We identify presence and fully characterize a previous unknown Wybutosine intermediate lacking a single methylation in the side chain (yW-14), the last step of the known biosynthetic pathway. Finally, from the list of known, fully characterized *S. cerevisiae* tRNA sequences, we use IMA to detect these known sequences from the pool of total tRNA.

## MATERIALS AND METHODS

### Transfer Ribonucleic Acid

Specific tRNA^**PHE**^ was purchased from Sigma-Aldrich (St Louis, MO.). Total tRNA from yeast was purchased from Invitrogen (Waltham, MA.). Specific tRNA^**PHE**^ was diluted to ∼ 400 ng/uL using DNase/RNase free water from Invitrogen (Waltham, MA.). Total tRNA was used without further processing.

### Liquid Chromatography

All separations were accomplished by reversed-phase liquid chromatography using an DNAPac™ RP, 4 μm, 2.1 mm X 100 mm column, (Thermo Fisher Scientific, CA.) on a Thermo Scientific™ Vanquish™ Horizon Quaternary UHPLC system (Thermo Fisher Scientific, San Jose, CA.). Mobile phase A consisted of 15mM dibutyl amine (Thermo Fisher Scientific, CA.), 25mM hexafluoro-2-propanol (Thermo Fisher Scientific, CA.). Mobile phase B consisted of 100% acetonitrile. Elution gradient of 10% B (from 0 to 0.5 min), 40% B at 7.0 min, 90% B at 7.1 held for 0.5 min, returning to 10% B at 7.51 min at a flow rate of 400 μL min^-1^. The column temperature was set at 80 °C.

### Mass Spectrometry

High resolution accurate mass analysis was performed on an Thermo Scientific™ Orbitrap Ascend™ Tribrid™ mass spectrometer (Thermo Fisher Scientific, CA.) interfaced with a heated electrospray (H-ESI) source in negative polarity mode. Global instrument conditions were sheath gas, auxiliary gas, and sweep gas of 50, 10 and 1 arbitrary units, respectively; ion transfer tube temperature of 350 °C; vaporizer temperature of 320 °C; and spray voltage of 3500 V.

Full scan acquisition was acquired in Intact Protein mode with low pressure setting, using a resolution of 240,000 at 400 m/z, mass range 800-3000 m/z, automatic gain control (AGC) 4e5, and max injection time (IT) 500 ms; quadrupole isolation of 1 m/z; and radio frequency (RF) of 50%; Data was acquired using Xcalibur™ 4.5 and analyzed with Freestyle™ 1.8 and BioPharma Finder™ 5.1.

Tandem Mass Spectrometry Acquisition was acquired in Peptide mode with low pressure setting, using a resolution of 60,000 at 400 m/z, mass range 500-2000 m/z, automatic gain control (AGC) 4e6, max injection time (IT) 200 ms and RF of 35%. Filters used were, Intensity threshold (2.5e4); Dynamic Exclusion 6 sec with both mass tolerances set to 5 ppm. Fragmentation was generated using CID and detected in the orbitrap at a resolution of 30,000. Ion trap settings were 35% CID energy, activation time of 10 ms, Q set to 0.25. Fragmentation scan range was set at 300-2000 Da with an injection time of 200 ms and a Normalized AGC target of 300%.

Deconvolution was performed using Thermo Scientific BioPharma Finder™ software 5.1, *Intact Mass Analysis*, using the Xtract deconvolution algorithm (Supplemental Figure 4). Chromatogram Trace Type was TIC, and *m/z* Range of 800 to 3000. Source Spectra Method was Averaged Over Selected Time. Deconvolution Algorithm was Xtract with an Output Mass Range of 24000 to 26000 with a S/N Threshold of 3 and a Rel. Abundance Threshold (%) of 3. Charge Range was set from 10 to 35 with a Min. Num Detected Charge of 5 using the Nucleotide Table and Negative Charge. Under Identification, Sequence Matching Mass Tolerance was set to 10 ppm with Multiconsensus Component Merge Mass Tolerance at 10 ppm. All tRNA sequences for PHE sample (Supplemental table 2) were created under Sequence Manager lacking the 3’ CCA motif and ending in a terminal phosphate.

### Nuclease Digestion

Nucleoside and oligonucleotide digests were performed as previously described.^38^ Briefly, an aliquot of tRNA was heated at 95°C for five minutes and cooled in a water bath. To the cooled sample was added 0.1U of P1 nuclease 0.1U acid phosphatase (Sigma Aldrich, St. Louis, MO.), and 0.01U RNase A (Worthington Biochemical), followed by heated at 37°C for two hours. Sample was removed, taken to dryness in a speed vac, resuspended in mobile phase A and injected onto a Accucore C18+ column 2.1x100mm, 1.5um (Thermo Fisher Scientific, CA.). Gradient conditions start at 0% B (from 0 to 3 min.), 25% B at 10 min., 99% B at 14 min. held for 1 min. then returning to 0% B at 15.1 min.

For oligonucleotide analysis, both tRNA^PHE^ and total tRNA were digested in RNase/DNase free water made 200 mM with ammonium acetate(Invitrogen, CA.), followed by introduction of 50U RNase T1 (Thermo Fisher Scientific, CA)/ug of tRNA. Sample was placed in a heat block and allowed to digest for two hours at 40°C, taken to dryness in a speed vac, resuspended in MPA and injected onto a Accucore C18+ column 2.1x100mm, 1.5um (Thermo Fisher Scientific, CA.) using 15mM DBA 25mM HFIP adjusted to pH 9.0 with ammonium acetate (Thermo Fisher Scientific, CA.) for Mobile Phase A, and 100% Methanol (Thermo Fisher Scientific, CA) for Mobile Phase B. Gradient conditions start at 10% B (from 0 to 1 min.), 30% B at 40 min., 35% B at 68 min. then increasing at 68.1 min to 90% B and held for 1 min. then returning to 10% B for 10 min.

## RESULTS

### UHPLC-HRAM-MS of tRNA^PHE^

Intact mass measurement of tRNA^PHE^ shows a charge state distribution between -9 to -26 with the most abundant charge states between -15 and -19 (**Figure 1A**). For each charge state several features are observed with a minimum and maximum relative abundance between 3 and 30 (Figure 1B). Deconvolution^46–48^ of the spectra from the main chromatographic peak returns 8 masses, the most abundant having a monoisotopic mass of 24610.450 Da. Two peaks at 30% relative abundance flanking the most abundant peak had deconvoluted masses at 24626.526 and 24596.547, a difference of +16.076 and -13.903 Da from the most abundant peak’s mass. Using the verified sequences^49^ of the two published isodecoders returns theoretical monoisotopic masses of 24859.521 and 24874.532 Da^50^ for a difference between the two published sequences of 16 Da and from the observed masses by 249.071 Da. The calculated masses include post-transcriptional modifications as well as the uncharged 3’ hydroxyl CCA motif on the acceptor stem. The difference of 249 Da is explained through the loss of 3’ adenosine in tRNA^PHE^ previously reported utilizing RNase T1 hydrolysis followed by anion exchange chromatography,^29^ later verified by Huang using intact mass spectrometry.^51^ By using the previous identification of truncation at the 3’ adenosine, we identify both published isodecoders, the predominately expressed tRNA^PHE^ having an adenosine at position 67, and the minor variant with a guanosine at position 67 complemented with cytosine at position 6. To identify other possible truncated species from the list of deconvoluted masses we constructed a series of masses to match against the deconvolution results.

**Figure 1:**
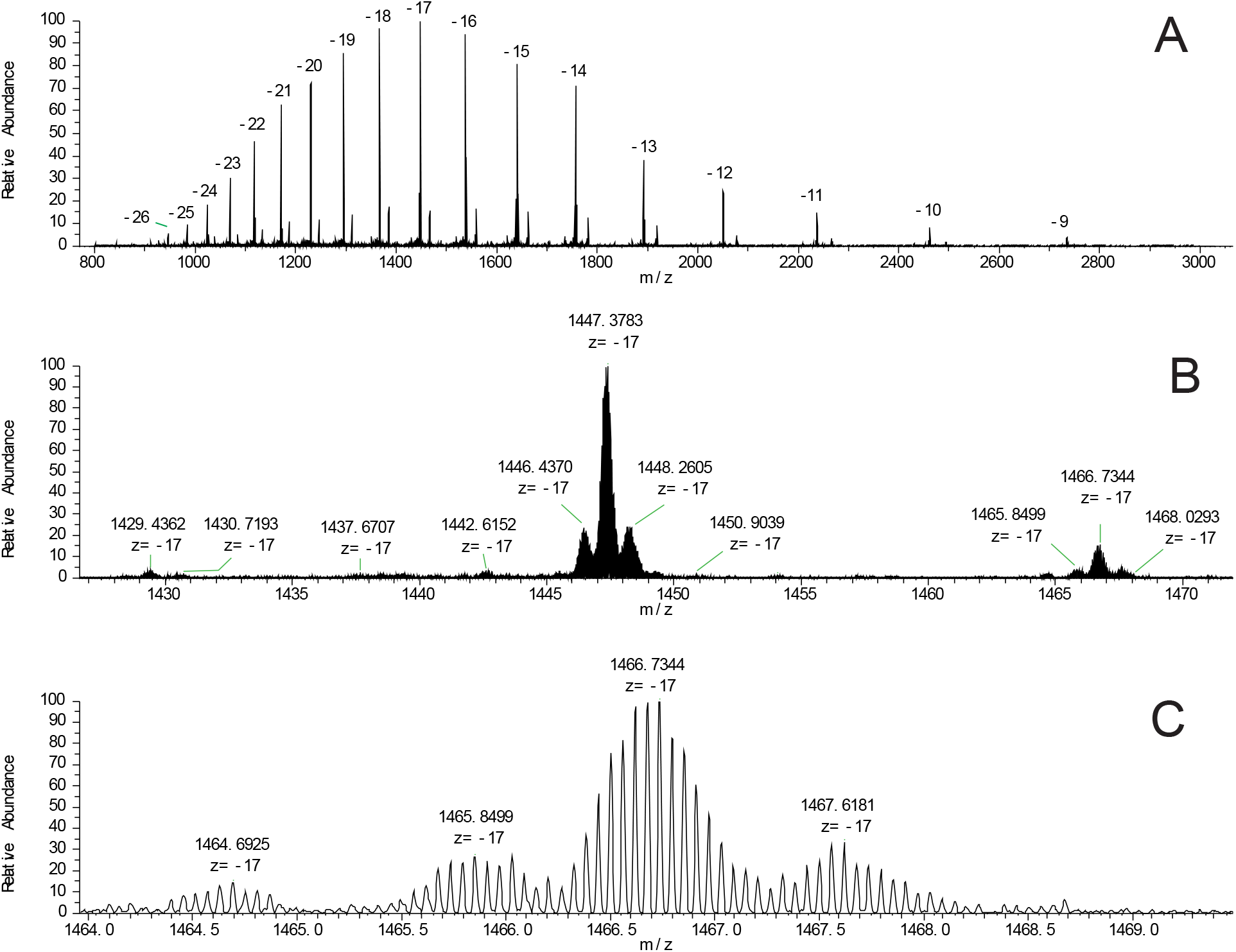
Intact mass spectrum of a commercially available tRNA^PHE^ standard. A: Charge state envelope of intact tRNA from -26 to -9. B: Zoomed image of baseline of highest abundant peak. C: Baseline isotopic resolution of the -17 charge state aids in deconvolution, and accurately identifies both isodecoders of tRNA^PHE^ as well as an A68 variant less one methylation.

A truncation series of monoisotopic intact masses for the A68 mature sequence was created with the largest mass being the full sequence with the 3’ terminal CpCpA and phosphorylated at the 5’ end. Each sequential mass removes either the base or the phosphate to generate a mass list of sequentially truncated sequences. When the calculated values were compared against the deconvoluted results, we were able to assign 8 out of 10 deconvoluted masses with the theoretical masses, with the most abundant (100% relative abundance) having the CpC truncation and CpCpA at ∼17% relative abundance.

### Identification of contaminating tRNAs

To identify the unknown deconvoluted masses, we performed UHPLC-HRAM-MS/MS nucleoside analysis^38^ to try to determine the presence of any contaminating tRNAs in the sample. Nucleoside analysis resulted in the detection of 24 modifications (**Supplemental Table 1**), and showed the presence of inosine, i^6^A, t^6^A, mcm^5^U, mcm^5^s^2^U, the rRNA modification m^6,6^A as well as m^2,2,7^G, a modification associated with miRNAs, as well as OHyW. *S. cerevisiae* lacks the enzyme TyW5^52^ necessary for its production of OHyW. Its presence is due to oxidation during the electrospray process, as the modifications elution time is the same as yW. The modifications m^6,6^A and m^2,2,7^G are attributed to contaminating RNAs. The isopentenyladenosine modifications, i^6^A and t^6^A, are found at position 37 of Wobble uridine anticodons. This reduces the number of contaminating unknowns down to 16, and when considering the two U34 modifications mcm^5^U and mcm^5^s^2^U, the number of possibilities is reduced to five, having both the detected position 34 and t^6^A modifications, tRNA^ARG^, tRNA^GLY^, tRNA^GLN^, tRNA^LYS^, and tRNA^GLU^.^53^ *S. cerevisiae* has 275 tRNA genes decoding the standard 20 amino acids,^54,55^ 35 of which have verified sequences tabulated in the Leipzig database with modifications within each tRNA’s sequence context. Searching these sequences for the detected modifications adds tRNA^VAL^ and tRNA^THR^ both having an inosine at position 34. Two other tRNA, tRNA^TYR^ (i^6^A at pos37), and tRNA^SER^ have both inosine and i^6^A, however the tRNA^SER^ have lengths that, when calculated, have masses outside the detected deconvoluted mass range (24000 – 26000 Da) and are therefore excluded from further consideration.

A truncation series was created for the remaining tRNA sequences containing i^6^, t^6^A and mcm^5^s^2^U. (**Supplemental Table 2**). Mass matching identified tRNA^LYS^ (3UU), tRNA^TYR^ (GUA), and tRNA^VAL^ (AAC). No other masses could be matched to the theoretical table. All identified sequences had a Matched Mass Error of < 6 ppm. Signature digestion products (SDP)^40^ for all non PHE decoders were then generated *in silico* as targets to search for in a full enzymatic digestion.

### Verification of contaminating tRNA identity

To verify the identity of the annotated tRNAs from the deconvolution experiments, enzymatic digestion was performed on the tRNA^PHE^ sample. SDPs for the tRNA sequence identified through deconvolution are shown in **Supplemental Table 3. Figure 2A** shows the MS/MS spectra of the tyrosine decoding tRNA signature digestion product. C and Y type ions are fully annotated and follow the predicted fragmentation pattern associated with RNA fragmentation^56^ with the C_3_ ion carrying the pos37 modification i^6^A. Tandem mass spectra for tRNA^LYS^ and tRNA^VAL^ were also found verifying presence in the sample (**Supplemental Figures 1 & 2**).

**Figure 2.**
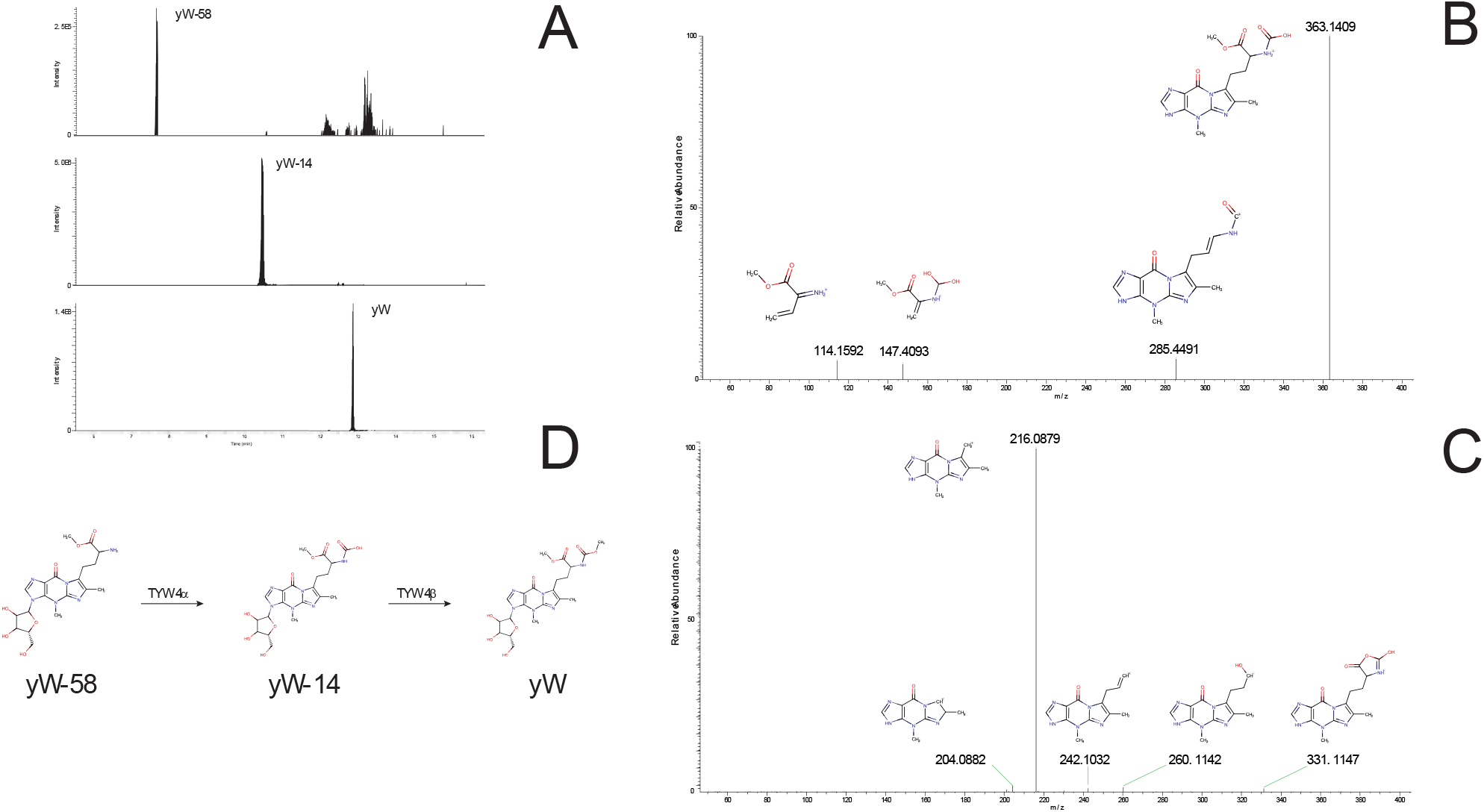
**A:** Extracted ion chromatograms for Wybutosine and its two immediate precursors yW-58 and yW-14. **B:** CID fragmentation spectra of the yW-14 modification. **C:** HCD fragmentation of the yW-14 modification, both HCD and CID fragment identification was performed using Fragment Identification Search. **D:** The predicted biosynthetic pathway, showing the yW-14 intermediate generated by the TYW4 enzyme during carboxymethylation.

### Identification of Wybutosine intermediate yW-14

One of the most abundant signals in the deconvolution results was identified as a demethylation of the A68 tRNA. Looking at the most probable labile methylation suggests a loss of methylation on the Wybutosine sidechain, which would result in a monoisotopic mass of 494.1761 Da. Extensive characterization of the yW biosynthetic pathway has previously been reported by Homa, et. al,^3^ with the last step in the enzymatic pathway a carboxymethylation of the yW-58 intermediate to yW. To verify if the demethylated signal in the deconvolution result was the yW-14 intermediate, extracted ion chromatogram (EIC) in the nucleoside data for the protonated mass were generated and returned a feature at ∼10.6 min, between yW-58 and yW (**Figure 3A**). Examining the collisional induced dissociation (CID) and High Energy Collision Dissociation (HCD) MS/MS filters for 495 (Figure 3B and 3C) yielded fragmentation consistent with a yW-14 structure. This suggests that during carboxymethylation from yW-58 to yW (**Figure 3D**), for the strain and conditions used to produce this commercial standard, the turnover from yW-14 to yW is slow enough to produce a measurable amount of intermediate tRNA detected in our deconvolution results.

**Figure 3.**
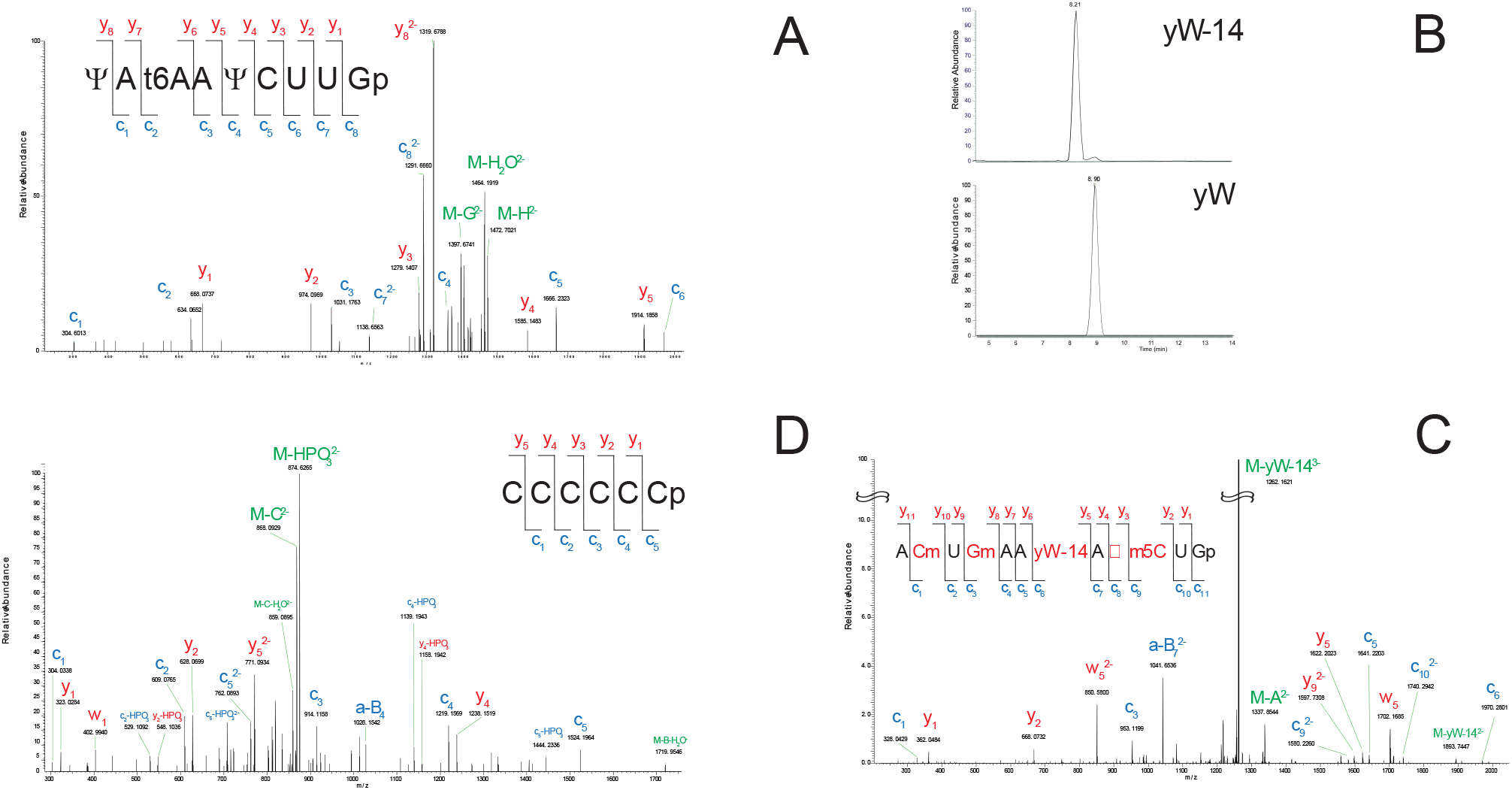
**A:** Signature digestion product from tRNA^TYR^. Insert shows fragmentation ladder. Fragment C3 carries the t6A modification from position 37 in the anticodon. **B**: Extracted ion chromatograms of the signature digestion products for the yW-14 and yW containing modification and yW. The oligonucleotide containing the yW-14 modification elutes before the oligonucleotide containing the yW modification. This elution profile would be expected for similar oligonucleotides differing by a methyl group. **C**: Tandem mass spectra for the oligonucleotide containing the yW-14 modification. Primary fragmentation channel is nucleobase loss of the tricyclic modification. Low abundant diagnostic fragment ions as well as internal canonical base loss confirms identity of oligonucleotide. **D**: 3’ terminal of proline decoding tRNA showing no adenosine terminus.

**Figure 4.**
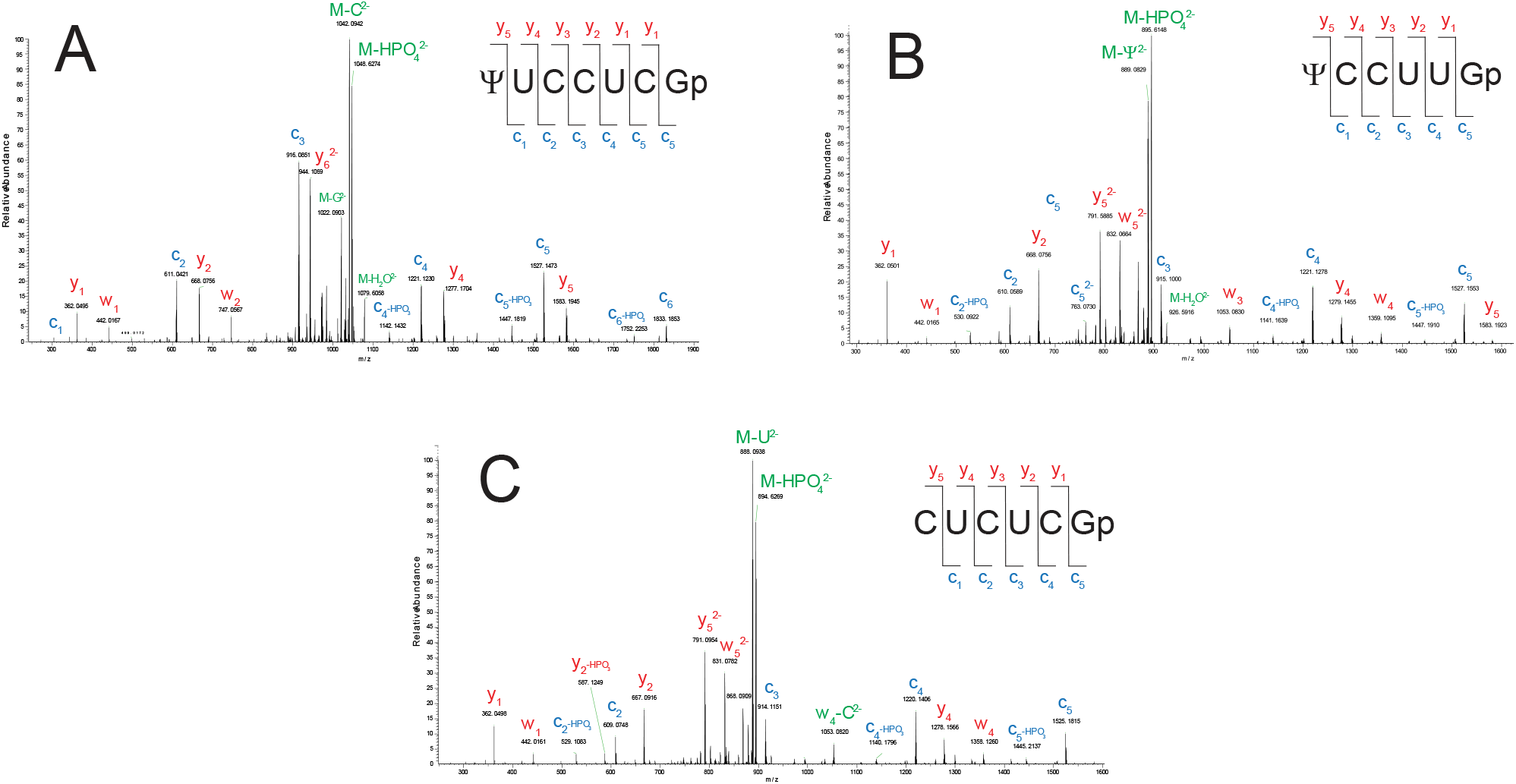
5’ terminal digestion products from A) tRNA^ARG^ICG, B) tRNA^LYS^ 3UU and C) tRNA^TYR^ GΨA.

The presence of yW-14 within the intact tRNA was then verified through oligonucleotide mass mapping. Using the online calculator Mongo Oligo v2.06 (http://rna.rega.kuleuven.be/masspec/mongo.htm) the verified sequence of tRNA^PHE^ was subjected to an *in silico* T1 digestion to generate the wybutosine containing signature digestion product with M = 4164.664 Da. A second *in silico* digestion was performed substituting yW for the yW-14 modification to generate a second SDP with M = 4151.656 Da for a difference of 13.008 Da. A full RNase T1 digestion of the tRNA^PHE^ standard was then performed and extracted ion chromatograms for the -2 and -3 charge state were generated. Both the yW and yW-14 containing oligonucleotides were detected in the -3 charge state with elution profiles differing by ∼ 40 seconds (**Figure 2B**). A difference in elution profile is expected for similar sequences differing only by a single methylation. Examining the spectra of both oligonucleotide shows primarily loss of the Wybutosine side chain. Low abundant diagnostic fragments are present and are used to verify sequence. During CID, oligonucleotides lose the terminal nucleotides sequentially. This allows for precise placement of the modified base within its sequence context. Additionally, due to sequential fragmentation, if one or more expected fragment ions are not observed in the spectra, yet others are, the observed sequence fragments remain diagnostic as their observed masses are dependent on the fragment masses before and after the observed fragment ion in the sequence. Figure 3C shows the yW-14 containing SDP with identified diagnostic fragment ions. Fragmentation profiles for both yW-14 and yW (**Supplemental Figure 3**) are, as expected, similar, with higher mass doubly and singly charged masses having a delta mass of 14 Da between the two oligonucleotides. Having identified the yW-14 variant in the sample we confidently assign the final unknown from the intact mass deconvolution results (**Table 1**).

**Table 1.** Intact mass deconvolution results.

| Modification | Monoisotopic Mass | Matched Mass Error (ppm) | Relative Abundance | Number of Charge States | Charge State Distribution |
| --- | --- | --- | --- | --- | --- |
| PHE A68 CpC | 24610.397 | 3.3 | 100.00 | 20 | 9-28 |
| PHE G68 CpC | 24626.384 | 3.3 | 25.77 | 19 | 9-27 |
| PHE A68 CpCp undermethylated yW | 24596.388 | 4.6 | 22.74 | 19 | 9.27 |
| PHE A68 5'p-CpCpA | 24939.448 | 3.1 | 18.55 | 18 | 10-27 |
| PHE G68 5'p-CpCpA | 24955.453 | 3.8 | 4.72 | 15 | 12-26 |
| TYR GUA CpC* | 24925.423 | 1.0 | 3.89 | 15 | 12-26 |
| PHE A68 C | 24305.494 | 3.3 | 3.23 | 14 | 13-26 |
| PHE A68 CpC | 24530.568 | 3.4 | 3.12 | 14 | 12-25 |

### Identification of tRNAs in a total tRNA pool by intact mass analysis

We next sought to apply the intact mass analysis to a pool of tRNA to see if it is possible to identify intact tRNA from total tRNA, and to detect if the 3’ adenosine truncation is specific for the tRNA^PHE^ sample. Using the same instrument parameters and chromatographic conditions an injection of *S. cerevisiae* total tRNA was acquired. Using a sliding window^57^ approach on the main chromatographic peak ninety-five masses were generated and matched against theoretical truncated masses to identify forty one identities, including the main phenylalanine decoding tRNA (**Table 2**). Seventeen of the identities contained the 3’ motif CpC and were predominately the most abundant masses. Only eight masses matched for the expected full sequence of tRNA with twelve tRNA matched masses differing by 1 Da. The 1 Da difference between calculated and observed is due to isotopomer envelope overlap from the A0 peak from one tRNA identity with that of the a second tRNA isotopomer envelope. ^48,58–60^ Continued improvements in chromatographic substrates as well as deconvolution algorithms will offset the isotopomer overlap allowing deconvolution without the 1 Da difference.

**Table 2.**
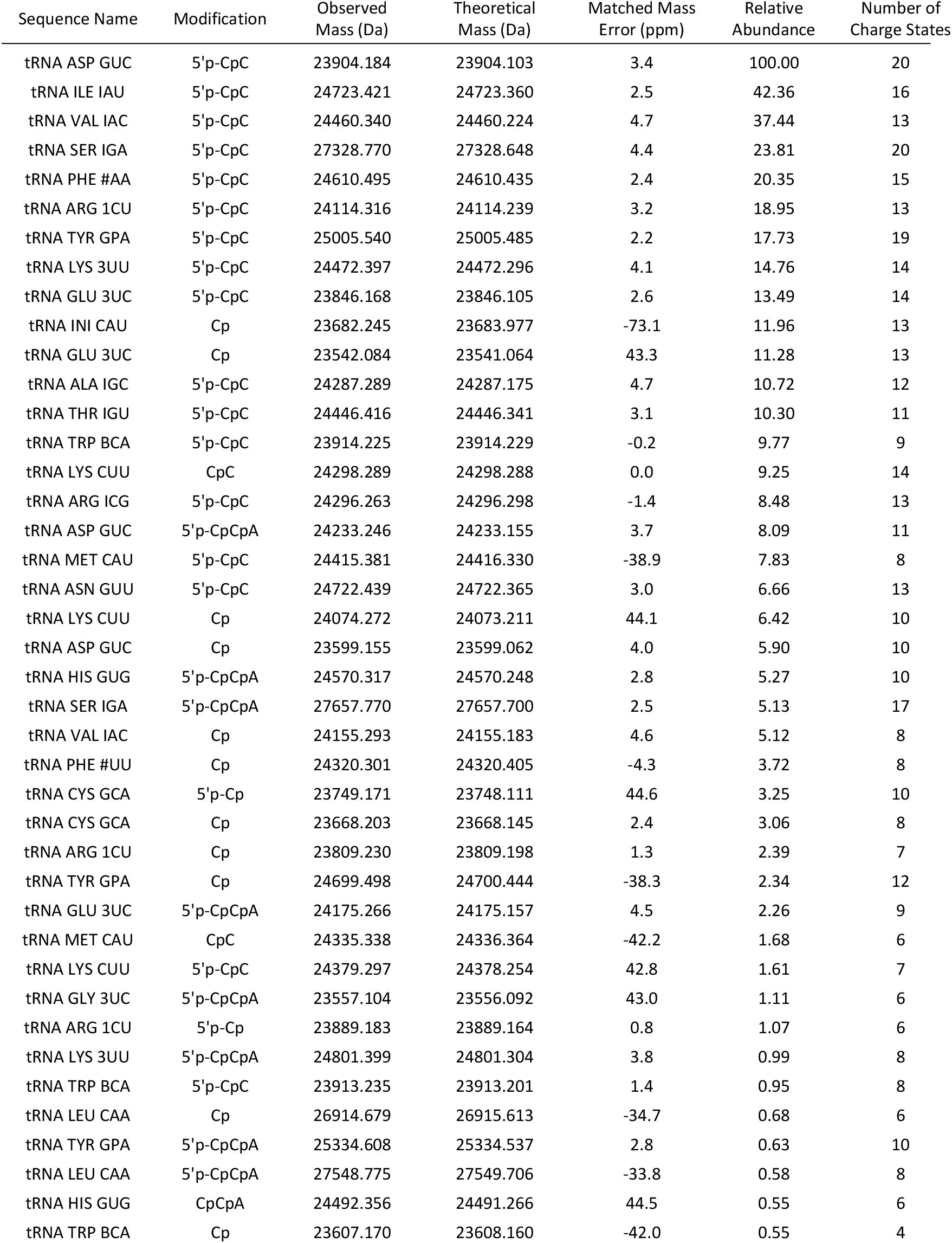
Deconvolution results from an acquisition of total tRNA from S. cerevisiae.

### Verification of 5’ and 3’ termini through oligonucleotide mass mapping

Verification of truncated tRNA acceptor stems was performed by mass mapping experiments using full enzymatic digestions, targeting the tRNAs containing the CpC motif as these signals were detected at the highest abundance. Reviewing the table of known sequences, we chose the 3’ termini of tRNA^PRO^ as it contains 6 sequential cytidine nucleotides and the 5’ termini of tRNA^ARG^. The arginine decoder was chosen as it also contains six nucleotides to increase specificity in identification by reducing redundancy.

(**Supplemental Table 4**). Additionally, the use of RNase T1 generates a 3’ guanosine phosphate during enzymatic digestion and is detected in the spectra at a *m/z* value of 362.00. Identification of the 3’ six nucleotide sequence CCCCCC of the tRNA^PRO^ isodecoder generated through a T1 digestion of *S. Cerevisiae* total tRNA is shown in **Figure 3D**. This oligonucleotide lacks the terminal adenosine and instead contains a 3’ terminal phosphate. Signature loss of phosphate from the precursor is detected as a doubly charged species and is the primary ion in the spectra. All c ions generated through CID are detected for the oligonucleotide. Secondary fragmentation, such as phosphate/water loss, is also detected for this oligonucleotide. Detection of the secondary fragments are due to the ionization efficiency of cytidine. Further digestions products lacking the 3’ termini were also identified (**Supplemental Figures 4-7**). Next, we sought to verify 5’ phosphorylation from the same digestion. *S cerevisiae* tRNA^ARG^ (ACG), tRNA^LYS^ (UUU), and tRNA^TYR^ (GUA) are the only tRNA in an RNase T1 digest whose 5’ terminal digestion product contains more than four nucleotides. RNase T1 *in silico* digestion product masses with and without 5’ phosphate was used to create extracted ion chromatograms. All three 5’ terminal oligonucleotides were found to not contain the 5’ phosphate (**Figure 4**). No masses were found which contained the 5’ terminal phosphate.

## DISCUSSION

Transfer RNA contain the greatest density of modifications than any other RNA. To date, over 130+ modifications have been characterized with more being discovered anew. These modifications are added post-transcriptionally and can range between a simple methylation on the nucleobase and/or the ribose, to extensively modified versions of their canonical requiring multiple enzymatic steps and cofactors. The biological implications of many modifications are still being uncovered; however, it is without question that modification of the nucleotide affects tRNA structure and function. One of the challenges associated with determining biological function of post transcriptional modifications is determining sequence placement of the modification. Next-generation sequencing (NGS) such as Illumina and Oxford nanopore are a promising technology for mapping tRNA modification within their respective sequence context.

One of the limitations of current NGS technology is its ability to discriminate between positional isomers or recognize hypomodifications as we have shown here with yW-14. To obtain this level of granularity in RNA modification mapping the use of mass spectrometry is needed. Studying tRNA by LC-MS/MS requires enzymatic digestion followed by two analyses. The first digestion reduces the intact RNA to the nucleoside level, followed by LC-MS/MS yielding a census of the modifications present in the sample. A second digestion is then performed using a base specific nuclease, to produce oligonucleotides of varying lengths which are separated chromatographically and analyzed using CID for accurate placement of the modification within its sequence context.

To analyze RNA by LC hyphenated mass spectrometry the use of ion pairing reagents are used to mask the electronegative backbone of the strand, increasing the biopolymer’s hydrophobic character to retain onto chromatographic substrates, such as C18. Triethyl amine (TEA) and hexafluorisopropylalcohol (HFIP) are common ion paring reagents, however the tertiary amine can only act as a hydrogen bond acceptor. The secondary structure of RNA is maintained through hydrogen bonding between nucleobase and their complement facilitated by atoms within the nucleotide which can participate either as a hydrogen bond donor or acceptor. The ratio of acceptor to donor sites is roughly 2:1 for canonical RNA nucleotides. By using a primary or secondary amine, such as dibutylamine, the number of hydrogen bond sites are exploited to increase the hydrophobic nature of the RNA. When the chromatographic substrate is inert, such as divinylbenzene, progressively longer nucleic acids can be retained without the need to pH the mobile phase, further increasing ionization. During the electrospray process the additional hydrophobicity assists the nucleic acid to desorb from the aqueous droplet more rapidly while simultaneously displacing metal adduction (Na^+^, K^+^, etc.) that reduce the signal of the RNA. The increase in desorption is witnessed by the shift to lower charge states becoming more abundant and carrying the amine as an H-bonded adduct. By applying in source fragmentation of ∼30-35 V the adduction is removed, and baseline isotopic peak envelopes with little to no metal adducts are observed.

The variation of the acceptor stem sequence is attributed to tRNA-nucleotidyltransferase, a template-independent RNA polymerase. ATP (CTP):tRNA-specific nucleotidyltransferase activity is required for tRNA biosynthesis and maturation.^61^ In yeast, the CCA motif is added onto the 3’ end of tRNA^62^ transcripts sequentially from their respective triphosphate precursors with the 3’ hydroxyl of the terminal adenosine complexing its specific amino acid. Using Intact Mass Analysis, we show that the tRNA exists in various forms of maturation, with the predominate species being CpC for all three variants. The abundance of the uncharged tRNA having the 3’ CpCpA was identified with mass deltas between theoretical and observed at < 4 ppm. Previous work show that deadenylation can happen under oxidative stress^63,64^ and could be a possible regulator of translation.^63^ To our knowledge, there has been no previous identification of tRNA containing more than the adenosine truncation at the 3’ termini, such as C or CpC by mass spectrometry. It is possible that the truncations could be the result of interruption of tRNA maturation homeostasis when the cells were harvested, however this doesn’t explain the abundance of the CpC termini. Examining published crystal structures for 3’ identification returns ambiguous results with some having a full 3’ CCA motif^65–68^ with others failing to show complete 3’ termini.^69–72^ The lack of phosphorylation for the 5’ terminal as well as verification of 3’ CpC through traditional mass mapping suggests that the process of acceptor stem maturation and 3’ aminoacylation is still not fully understood.

## CONCLUSIONS

Here we show the utility of high resolution accurate mass spectrometry coupled to an ion pairing chromatography platform using a hydrogen bond donor amine for the intact mass analysis of tRNA. The use of a donor amine increases the hydrophobicity of the RNA resulting in a faster ejection from the electrosprayed droplet while simultaneously suppressing metal adduction. This rapid desorption from the droplet results in a relatively clean charge state signal in the spectra which can be deconvoluted into the accurate mass. For the analysis of tRNAs this precision allows for facile discrimination between known sequences with sensitivities of single base substitutions or demethylations. The analytical approach presented here has for the first time shown the ability to identify individual tRNAs from an injection of total tRNA opening the possibility for the rapid screening of RNA and their modifications.

## DATA AVAILABILITY STATEMENT

The data underlying this article will be shared on reasonable request to the corresponding author.

## SUPPLEMENTARY DATA

Supplied separately.

## ACKNOWLEDGEMENTS

The authors would like to thank Hiroshi Nakayama (RIKEN, Japan), Vivian Cheung (University of Michigan), and Rafael Morin (Thermo Fisher Scientific) for reviewing the manuscript and providing critical feedback.

