## Supplemental Figures for "Comprehensive Characterization of tRNA by Ultra High-Performance Liquid Chromatography High-Resolution Accurate Mass Spectrometry"

### Supplemental Figure Legend

**Supplemental Figure 1**: Annotated spectra from signature digestion product for tRNA$^{LYS}$

**Supplemental Figure 2**: Annotated spectra from signature digestion product for tRNA$^{VAL}$

**Supplemental Figure 3**: Annotated spectra from signature digestion product for tRNA$^{PHE}$

**Supplemental Figure 4:** Annotated 3' digestion product for tRN$^{VAL}$ (CAC) presenting no terminal adenosine.

**Supplemental Figure 5:** Annotated 3' digestion product for tRN$^{VAL}$ (IAC) presenting no terminal adenosine.

**Supplemental Figure 6:** Annotated 3' digestion product for tRN$^{ILE}$ (IAU) presenting no terminal adenosine.

**Supplemental Figure 7:** Annotated 3' digestion product for tRN$^{ALA}$ (IGC) presenting no terminal adenosine.

**Supplemental Figure 1:** Annotated signature digestion product for tRNA $^{LYS}$. Fragment $C_3$ (1004 Da) helps in identification of the oligonucleotide. Dense fragmentation around 1600 Da due to side chain fragmentation of both mcm$^5$s$_2$U as well as t$^6$A (insert)

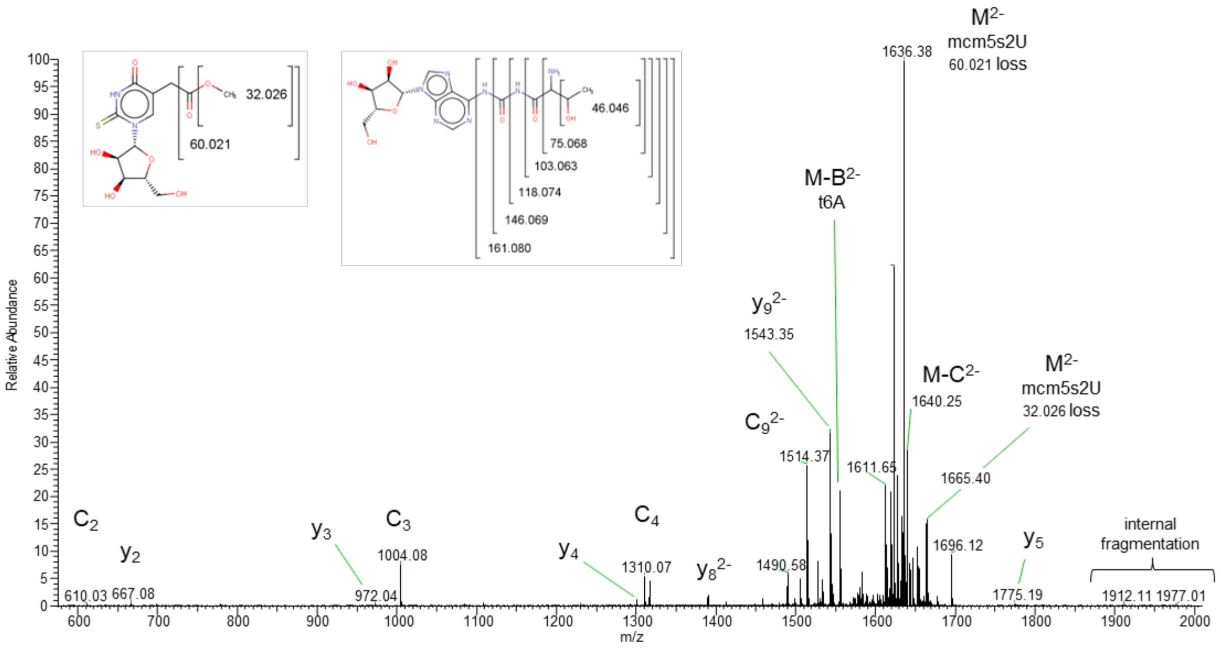

**Supplemental Figure 2:** Annotated spectra from $tRNA^{VAL}$ signature digestion product

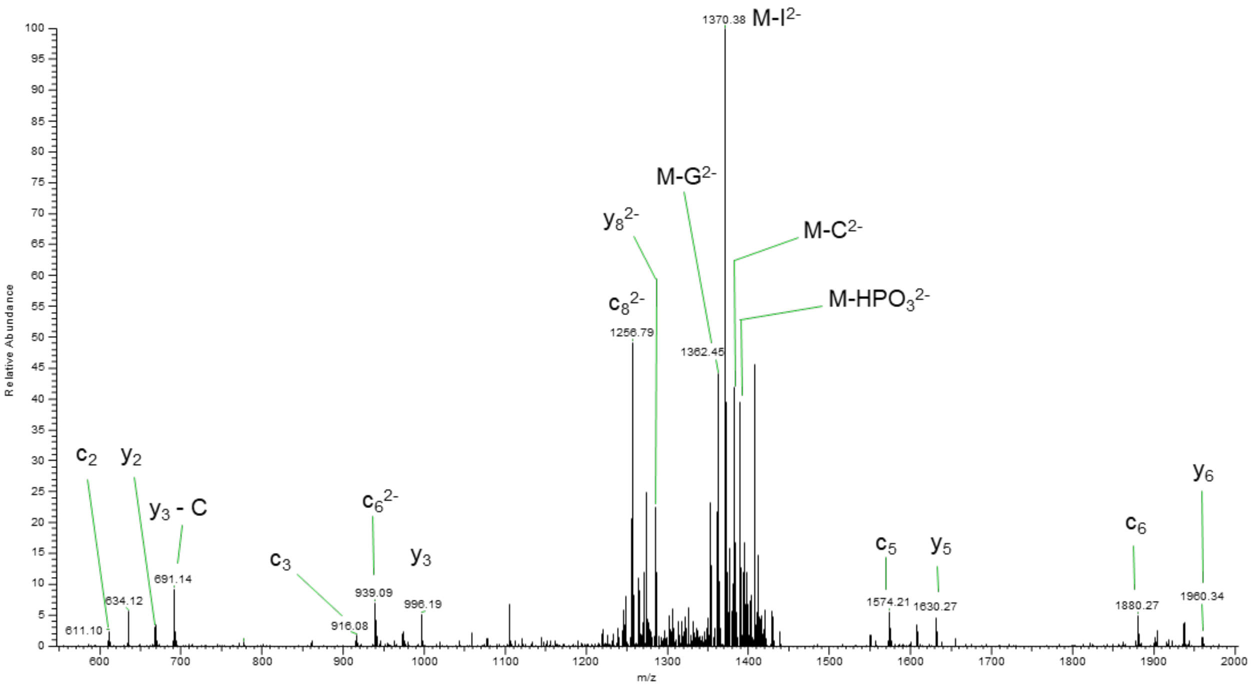

**Supplemental Figure 3:** Annotated signature digestion product for $tRNA^{PHE}$. Fragmentation of the yW containing oligonucleotide is primarily loss of the yW base.

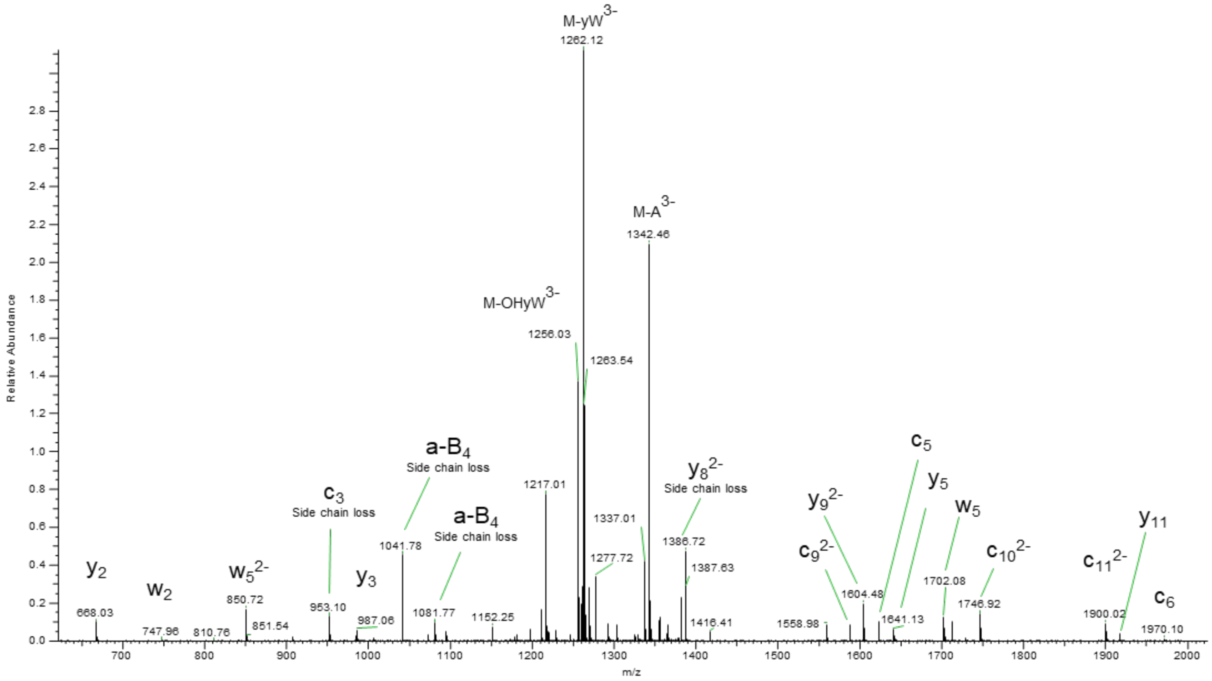

**Supplemental Figure 4:** Annotated 3’ digestion product for $tRN^{VAL}$ (CAC) presenting no terminal adenosine.

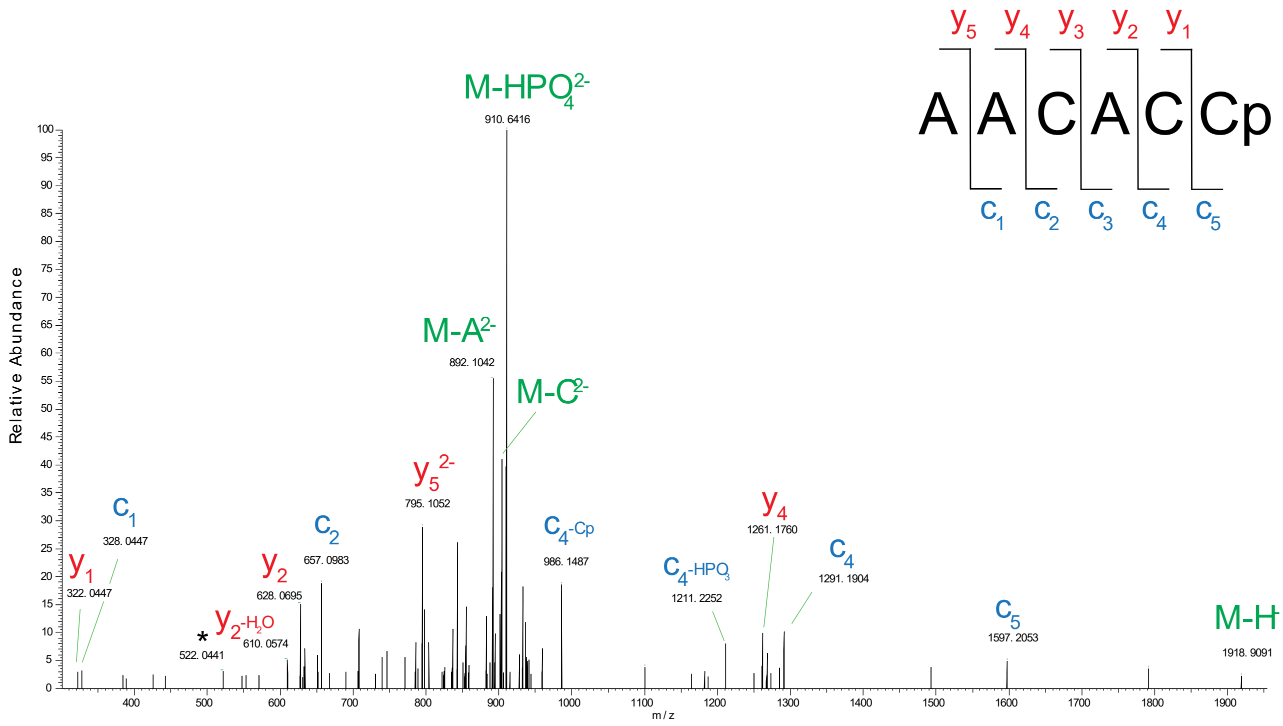

**Supplemental Figure 5:** Annotated 3’ digestion product for tRN$^{VAL}$ (IAC) presenting no terminal adenosine.

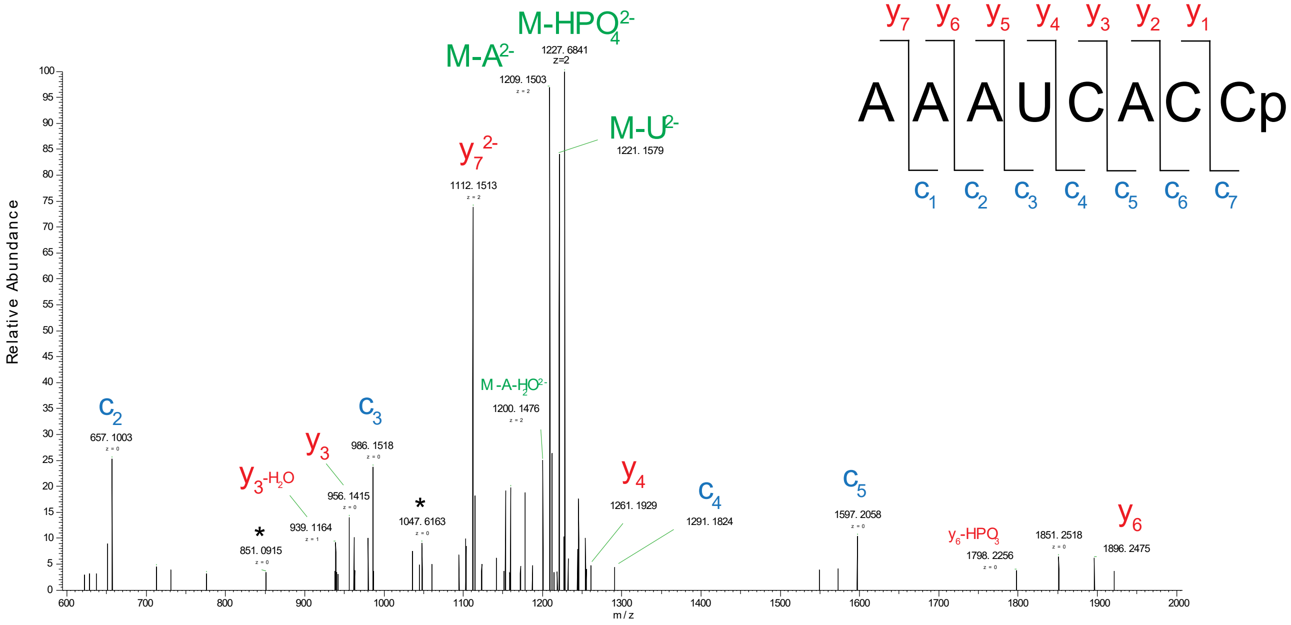

**Supplemental Figure 6:** Annotated 3' digestion product for tRN$^{ILE}$ (IAU) presenting no terminal adenosine.

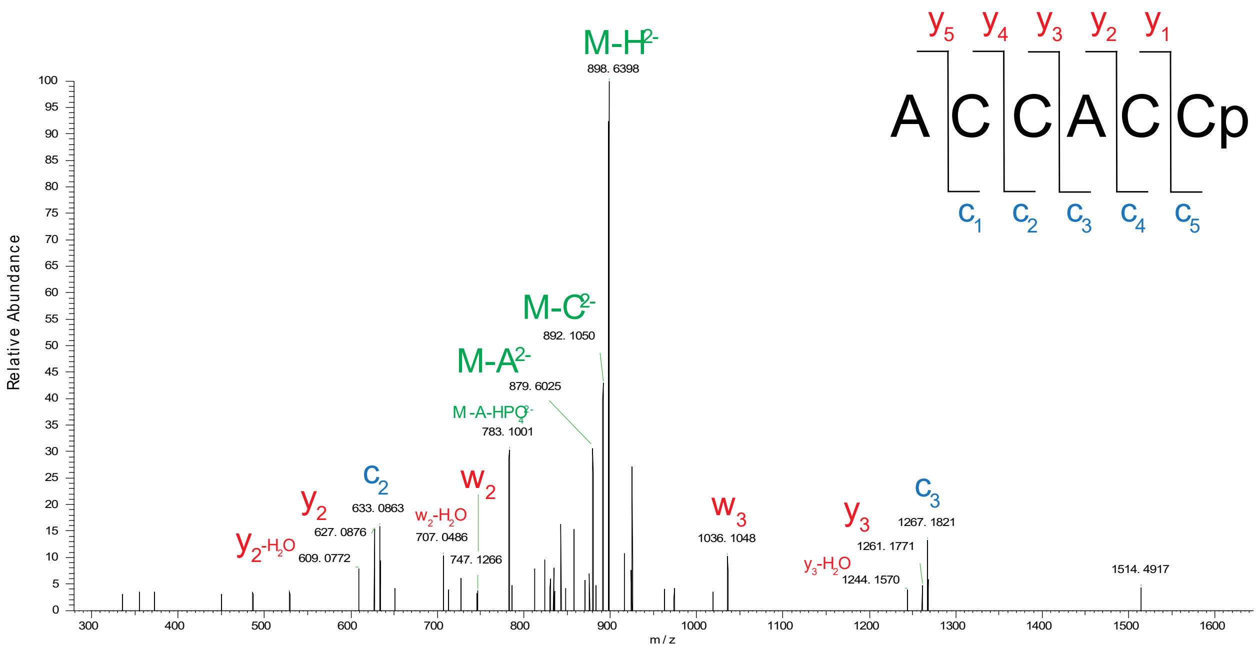

**Supplemental Figure 7:** Annotated 3' digestion product for tRN$^{ALA}$ (IGC) presenting no terminal adenosine.

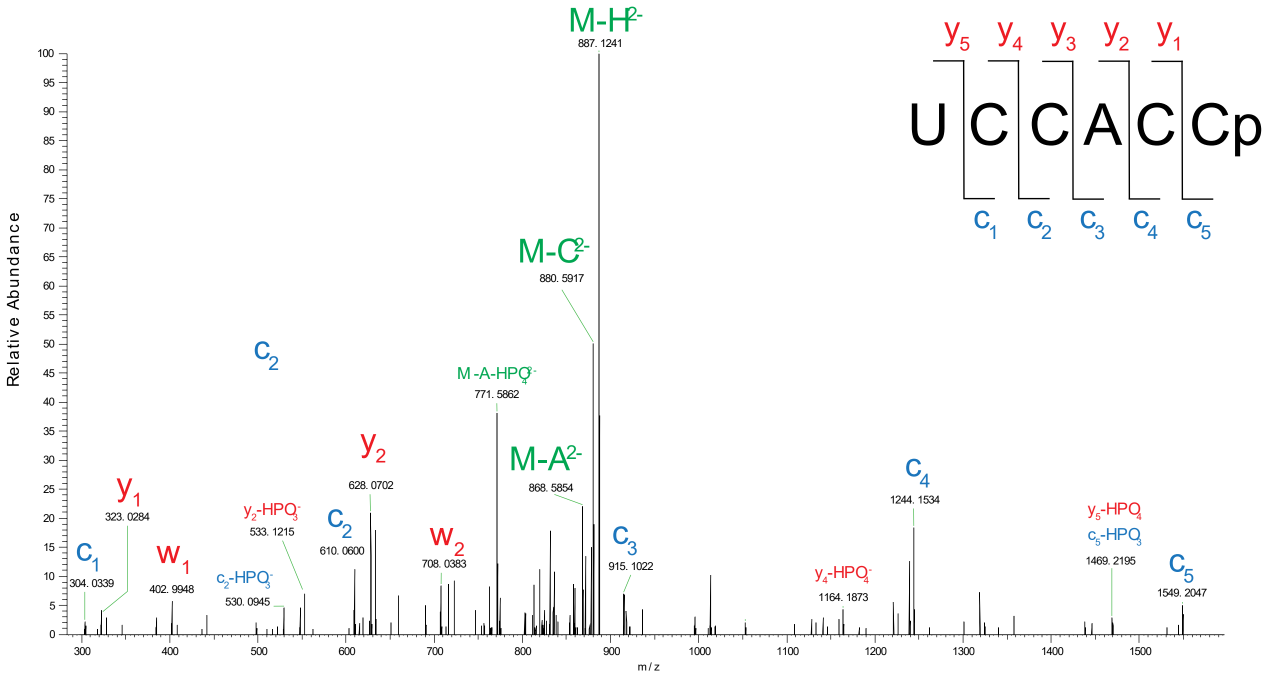
