## Supplementary Tables for "Comprehensive Characterization of tRNA by Ultra High-Performance Liquid Chromatography High-Resolution Accurate Mass Spectrometry"

**Supplemental Table Legend**

**Supplemental Table 1**: Nucleosides detected in tRNA$^{PHE}$ standard digest.

**Supplemental Table 2:** Verified sequences of *S. cerevisiae* containing modifications detected in nucleoside analysis and containing possible truncated 3' masses.

**Supplemental Table 3:** Signature digestion products for detected tRNA contaminates.

**Supplemental Table 4:** *In silico* generated 3' digestion products lacking the terminal adenosine.

**Supplemental Table 1:** Nucleosides detected in tRNA$^{PHE}$ standard digest.

| Name | Formula | DMass [ppm] | Calc. MW | RT [min] |
|---|---|---|---|---|
| D | C9 H14 N2 O6 | -1.06 | 246.08493 | 0.819 |
| P | C9 H12 N2 O6 | -0.47 | 244.06942 | 0.823 |
| m3C | C10 H15 N3 O5 | 0.04 | 257.10118 | 1.501 |
| m1A | C11 H15 N5 O4 | 0.23 | 281.11247 | 1.737 |
| m5C | C10 H15 N3 O5 | -0.39 | 257.10107 | 2.203 |
| Cm | C10 H15 N3 O5 | -0.65 | 257.101 | 2.698 |
| m7G | C11 H15 N5 O5 | -0.54 | 297.10716 | 2.769 |
| I | C10 H12 N4 O5 | -0.07 | 268.08075 | 2.892 |
| m5U | C10 H14 N2 O6 | -0.5 | 258.08506 | 3.482 |
| Gm | C11 H15 N5 O5 | -0.39 | 297.1072 | 5.988 |
| mcm5U | C12 H16 N2 O8 | -0.25 | 316.09058 | 6.304 |
| m2G | C11 H15 N5 O5 | -0.41 | 297.1072 | 6.385 |
| t6A | C15 H20 N6 O8 | -0.85 | 412.13391 | 6.648 |
| m2,2,7G | C13 H19 N5 O5 | -0.62 | 325.13842 | 7.493 |
| m2,2G | C12 H17 N5 O5 | -0.4 | 311.12284 | 7.685 |
| yw-58* | C19 H26 N6 O7 | -4.07 | 450.18447 | 7.687 |
| Am | C11 H15 N5 O4 | -0.41 | 281.11229 | 7.804 |
| mcm5s2U | C12 H16 N2 O7 S | 0.1 | 332.06786 | 8.239 |
| m6A | C11 H15 N5 O4 | -0.4 | 281.11229 | 8.7 |
| yw-86* | C17 H22 N6 O7 | 0.77 | 422.15532 | 9.754 |
| yw-14 | C20 H26 N6 O9 | -0.24 | 494.17601 | 10.482 |
| m6,6A | C12 H17 N5 O4 | -0.26 | 295.12798 | 11.447 |
| OHyW | C21 H28 N6 O10 | -0.08 | 524.18665 | 12.869 |
| yW | C21 H28 N6 O9 | -0.9 | 508.19132 | 12.87 |
| i6A | C15 H21 N5 O4 | -0.01 | 335.15935 | 13.679 |

* No MS/MS

**Supplemental Table 2**: Verified sequences of S. cerevisiae containing modifications detected in nucleoside analysis and containing possible truncated 3' masses.

<table>
<tr><th></th><th></th><th></th><th></th><th></th><th>AA</th><th>CpCpA-aa</th><th>5'p-CpCpA</th><th>CpCpA</th><th>CpCp</th><th>CpC</th><th>Cp</th></tr>
<tr><td>>tdbR00000012</td><td>Ala|IGC</td><td>GGGCGUGUKGCGUAGDCGGDAGCGCRCUCCCUUIGCOPGGGAGAGGDCUCCGGTPCGAUUCC<br>GGACUCGUCCACCA</td><td>M</td><td>=</td><td>71.03711</td><td>24590.29507</td><td>24616.227</td><td>24536.2607</td><td>24287.175</td><td>24207.209</td><td>23982.134</td></tr>
<tr><td>>tdbR00000370</td><td>Arg|1CU</td><td>GCUCGCGUKLCGUAADGGCAACGCRPCUGACU1CU6APCAGAAGADUAUGGGTPCG"CCCCCAU<br>CGUGAGUGCCA</td><td>M</td><td>=</td><td>156.10111</td><td>24502.42307</td><td>24443.291</td><td>24363.3247</td><td>24114.239</td><td>24034.273</td><td>23809.198</td></tr>
<tr><td>>tdbR00000371</td><td>Arg|1CU</td><td>GCUUGCGUKLCGUAADGGCAACGCRPCUGACU1CU6APCAGAAGADUAUGGGTPCG"CCCCCAU<br>CGUGAGUCCCA</td><td>M</td><td>=</td><td>156.10111</td><td>24463.40107</td><td>24404.269</td><td>24324.3027</td><td>24075.217</td><td>23995.251</td><td>23770.176</td></tr>
<tr><td>>tdbR00000369</td><td>Arg|ICG</td><td>PUCCUCGUKLCCCAADGGDCACGGCRPCUGGCUICGAACCAGAAGADU?CAGGTPCA"GUCCUG<br>GCGGGGAAGCCA</td><td>M</td><td>=</td><td>156.10111</td><td>24684.48207</td><td>24625.35</td><td>24545.3837</td><td>24296.298</td><td>24216.332</td><td>23991.257</td></tr>
<tr><td>>tdbR00000300</td><td>Asn|GUU</td><td>GACUCCAUGLCCAAGDDGGDDAAGGCRUGCGACUGUU6APCGCAAGAD?GUGAGTPCA"CCCUC<br>ACUGGGGUCGCCA</td><td>M</td><td>=</td><td>114.04293</td><td>25068.49089</td><td>25051.417</td><td>24971.4507</td><td>24722.365</td><td>24642.399</td><td>24417.324</td></tr>
<tr><td>>tdbR00000035</td><td>Asp|GUC</td><td>UCCGUGAUAGUUPAADGGDCAGAAUGGGCGCAUGUCKCGUGCCAGAU?GGGGTPCAAUUCCCC<br>GUCGCGGAGCCA</td><td>M</td><td>=</td><td>115.02694</td><td>24251.2129</td><td>24233.155</td><td>24153.1887</td><td>23904.103</td><td>23824.137</td><td>23599.062</td></tr>
<tr><td>>tdbR00000021</td><td>Cys|GCA</td><td>GCUCGUAUGGCGCAGDGGDAGCGCAGCAGAPUGCA+APCUGUUG7D?CUUAGTPCG"UCCUGAG<br>UGCGAGCUCCA</td><td>M</td><td>=</td><td>103.00919</td><td>24308.27815</td><td>24302.238</td><td>24222.2717</td><td>23973.186</td><td>23893.22</td><td>23668.145</td></tr>
<tr><td>>tdbR00000054</td><td>Glu|3UC</td><td>UCCGAUAUAGUGPAACGGCDAUCACAPCACGCU3UCACCGUGGAGAC?GGGGTPCGACUCCCC<br>GUAUCGGAGCCA</td><td>M</td><td>=</td><td>129.04259</td><td>24207.23055</td><td>24175.157</td><td>24095.1907</td><td>23846.105</td><td>23766.139</td><td>23541.064</td></tr>
<tr><td>>tdbR00000130</td><td>Gly|3CC</td><td>GGGCGGUUAGUGPAGDGGDDAUCAUCCCACCCU3CCAAGGUGGGGACACGGGTPCGAUUCUC<br>GUACCGCUCACCA</td><td>M</td><td>=</td><td>57.02146</td><td>24175.17742</td><td>24215.125</td><td>24135.1587</td><td>23886.073</td><td>23806.107</td><td>23581.032</td></tr>
<tr><td>>tdbR00000129</td><td>Gly|GCC</td><td>GCGBAAGUKGUUPAGDGGDAAAAUCCAACGPUGCCAPCGUUGGGC?CCGGTPCGAUUCCGGGC<br>UUGCGCACCA</td><td>M</td><td>=</td><td>57.02146</td><td>23516.14442</td><td>23556.092</td><td>23476.1257</td><td>23227.04</td><td>23147.074</td><td>22921.999</td></tr>
<tr><td>>tdbR00000144</td><td>His|GUG</td><td>GGCC:UCUUAGUAPAGD#GDDAGUACACAACAPUGUGKCAGUUGAAAC?CUGGTPCGAUUCUAG<br>GAGGUGGCACCA</td><td>M</td><td>=</td><td>137.05891</td><td>24611.32187</td><td>24571.232</td><td>24491.2657</td><td>24242.18</td><td>24162.214</td><td>23937.139</td></tr>
<tr><td>>tdbR00000145</td><td>His|GUG</td><td>GGCC:UCUUAGUAPAGD#GDDAGUACACAUCGPUGUGKCCGAUGAAAC?CUGGTPCGAUUCUAG<br>GAGAUGGCACCA</td><td>M</td><td>=</td><td>137.05891</td><td>24610.33787</td><td>24570.248</td><td>24490.2817</td><td>24241.196</td><td>24161.23</td><td>23936.155</td></tr>
<tr><td>>tdbR00000170</td><td>Ile|IAU</td><td>GGUCUCUUKLCCCAGDDGGDDAAGGCACCGUGCUIAUct6AACGCGGGGAD?AGCGGTPCG"UCC<br>CGCUAGAGACCACCA</td><td>M</td><td>=</td><td>113.08406</td><td>25051.44332</td><td>25035.328</td><td>24955.362</td><td>24706.276</td><td>24626.31</td><td>24401.235</td></tr>
<tr><td>>tdbR00000170</td><td>Ile|IAU</td><td>GGUCUCUUKLCCCAGDDGGDDAAGGCACCGUGCUIAU6ACGCGGGGAD?AGCGGTPCG"UCCCG<br>CUAGAGACCACCA</td><td>M</td><td>=</td><td>113.08406</td><td>25068.52702</td><td>25052.412</td><td>24972.4457</td><td>24723.36</td><td>24643.394</td><td>24418.319</td></tr>
<tr><td>>tdbR00000526</td><td>Ini|CAU</td><td>AGCGCCGUKLCGCAGDGGAAGCGCRCAGGGCUCAU6ACCCUGAU7D??UCGGAUCG"AACCG^G<br>CGGCGCUACCA</td><td>M</td><td>=</td><td></td><td>24534.45096</td><td>24631.42</td><td>24551.4537</td><td>24302.368</td><td>24222.402</td><td>23997.327</td></tr>
<tr><td>>tdbR00000249</td><td>Leu|CAA</td><td>GGUUGUUUGLCMGAGC#GDCDAAGGCRCCUGAPU?AAKCACAGGUAUCGUAAGAUG?AAGAGTP<br>CGAAUCUCUUAGCAACCACCA</td><td>M</td><td>=</td><td>113.08406</td><td>27565.82102</td><td>27549.706</td><td>27469.7397</td><td>27220.654</td><td>27140.688</td><td>26915.613</td></tr>
<tr><td>>tdbR00000251</td><td>Leu|UAA</td><td>GGAGGGUUGLCMGAGD#GDCDAAGGCRGCAGABUUAAKAPCUGUUGGACGGUUGUCCG?GCGA<br>GTPCG"ACCUCGCAUCCUUCACCA</td><td>M</td><td>=</td><td>113.08406</td><td>28310.93102</td><td>28294.816</td><td>28214.8497</td><td>27965.764</td><td>27885.798</td><td>27660.723</td></tr>
<tr><td>>tdbR00000250</td><td>Leu|UAG</td><td>GGGAGUUUGLCMGAGD#GDDDAAGGCRPCAGAPUUAGKCACUGAUAUCUUCGGAUG?AAGGGT<br>PCG"AUCCCUUAGCUCUCACCA</td><td>M</td><td>=</td><td>113.08406</td><td>27557.75502</td><td>27541.64</td><td>27461.6737</td><td>27212.588</td><td>27132.622</td><td>26907.547</td></tr>
<tr><td>>tdbR00000193</td><td>Lys|)UU</td><td>PCCUUGUUALCUCAGDDGGDAGAGCRPPCGGCU)UU6ACCGAAAU7D?AGGGGTPCG"GCCCCCU<br>APGAGGAGCCA</td><td>M</td><td>=</td><td>128.09496</td><td>24845.47892</td><td>24814.353</td><td>24734.3867</td><td>24485.301</td><td>24405.335</td><td>24180.26</td></tr>
<tr><td>>tdbR00000192</td><td>Lys|CUU</td><td>GCCUUGUULLCGCAADCGGDAGCGCRPAUGACUCUU6APCAUAAG7UUAGGGGTPCG"GCCCCC<br>UACAGGGCUCCA</td><td>M</td><td>=</td><td>128.09496</td><td>24738.43192</td><td>24707.306</td><td>24627.3397</td><td>24378.254</td><td>24298.288</td><td>24073.213</td></tr>
<tr><td>>tdbR00000284</td><td>Met|CAU</td><td>GCUUCAGUALCUCAGDAGGAAGAGCRPCAGPCUCAU6APCUGAAG7D?GAGAGTPCG"ACCUCUC<br>CUGGAGCACCA</td><td>M</td><td>=</td><td>131.04049</td><td>24779.45345</td><td>24745.382</td><td>24665.4157</td><td>24416.33</td><td>24336.364</td><td>24111.289</td></tr>
<tr><td>>tdbR00000284</td><td>Met|CAU</td><td>GCUUCAGUALCUCAGDAGGAAGAGCRPCAGPCUCAUc6APCUGAAG7D?GAGAGTPCG"ACCUCU<br>CCUGGAGCACCA</td><td>M</td><td>=</td><td>131.04049</td><td>24762.36975</td><td>24728.298</td><td>24648.332</td><td>24399.245</td><td>24319.279</td><td>24046.204</td></tr>
<tr><td>>tdbR00000083</td><td>Phe|#AA</td><td>GCGGAUUUALCUCAGDDGGGAGAGCRCCAGABU#AAYAP?UGGAG7UC?UGUGTPCG"UCCACAG<br>AAUUCGCACCA</td><td>M</td><td>=</td><td>147.06841</td><td>24989.58637</td><td>24939.487</td><td>24859.5207</td><td>24610.435</td><td>24530.469</td><td>24305.394</td></tr>
<tr><td>>tdbR00000084</td><td>Phe|#AA</td><td>GCGGACUUALCUCAGDDGGGAGAGCRCCAGABU#AAYAP?UGGAG7UC?UGUGTPCG"UCCACAG<br>AGUUCGCACCA</td><td>M</td><td>=</td><td>147.06841</td><td>25004.59737</td><td>24954.498</td><td>24874.5317</td><td>24625.446</td><td>24545.48</td><td>24320.405</td></tr>
<tr><td>>tdbR00000323</td><td>Pro|NGG</td><td>GGGBGUGUKGUCAAGDGGDAUGAUUCUCGCAUNGGKPGCGAGAG7CCCUGGGTPCA"UUCCCA<br>GCUCGCCCCCCA</td><td>M</td><td>=</td><td>97.05276</td><td>24109.21072</td><td>24109.127</td><td>24029.1607</td><td>23780.075</td><td>23700.109</td><td>23475.034</td></tr>
<tr><td>>tdbR00000324</td><td>Pro|NGG</td><td>GGGBGUGUGGUCAAGDGGDAUGAUUCUCGCAUNGGKPGCGAGAG7CCCUGGGTPCA"UUCCCA<br>GCUCGCCCCCCA</td><td>M</td><td>=</td><td>97.05276</td><td>24095.19472</td><td>24095.111</td><td>24015.1447</td><td>23766.059</td><td>23686.093</td><td>23461.018</td></tr>
<tr><td>>tdbR00000407</td><td>Ser|IGA</td><td>GGCAACUUGGCMGAGD#GDDAAGGCRAAAGAPUIGA+APCUUUJGGGCUCUGCCCG?GCAGGTP<br>CAAAUCCUGCAGUUGUCGCCA</td><td>M</td><td>=</td><td>87.03203</td><td>27614.78899</td><td>27624.726</td><td>27544.7597</td><td>27295.674</td><td>27215.708</td><td>26990.633</td></tr>
<tr><td>>tdbR00000408</td><td>Ser|IGA</td><td>GGCAACUUGGCMGAGD#GDDAAGGCRAAAGAPUIGA+APCUUUJGGGCUUUGCCCG?GCAGGTP<br>CGAGUCCUGCAGUUGUCGCCA</td><td>M</td><td>=</td><td>87.03203</td><td>27647.76299</td><td>27657.7</td><td>27577.7337</td><td>27328.648</td><td>27248.682</td><td>27023.607</td></tr>
<tr><td>>tdbR00000406</td><td>Ser|NGA</td><td>GGCACUAUGGCMGAGD#GDDAAGGCRACAGA<UNGA+APCUGUJGGGCUCUGCCCG?GCUGGT<br>PCAAAUCCUGCUGGUGUCGCCA</td><td>M</td><td>=</td><td>87.03203</td><td>27181.71199</td><td>27191.649</td><td>27111.6827</td><td>26862.597</td><td>26782.631</td><td>26557.556</td></tr>
<tr><td>>tdbR00000443</td><td>Thr|IGU</td><td>GCUUCUAUGLCCAAGDDGGDAAGGCRCCACA'UIGU6APGUGGAGAD?AUCGGTPCA"AUCCGAUU<br>GGAAGCACCA</td><td>M</td><td>=</td><td>101.04768</td><td>24779.47164</td><td>24775.393</td><td>24695.4267</td><td>24446.341</td><td>24366.375</td><td>24141.3</td></tr>
<tr><td>>tdbR00000444</td><td>Thr|IGU</td><td>GCUUCUAUGLCCAAGDDGGDAAGGCRCCACA'UIGU6APGUGGAGAD?GUCGGTPCA"AUCCGAC<br>UGGAAGCACCA</td><td>M</td><td>=</td><td>101.04768</td><td>24794.48264</td><td>24790.404</td><td>24710.4377</td><td>24461.352</td><td>24381.386</td><td>24156.311</td></tr>
<tr><td>>tdbR00000494</td><td>Trp|BCA</td><td>GAAGCGGUKLCUCAAD#GDAGAGCAPPCGABUBCAAAPCGAAGG7DUGCAGGTPCA"UUCCUGPC<br>CGUUUCACCA</td><td>M</td><td>=</td><td>186.07931</td><td>24331.36327</td><td>24242.253</td><td>24162.2867</td><td>23913.201</td><td>23833.235</td><td>23608.16</td></tr>
<tr><td>>tdbR00000555</td><td>Tyr|GPA</td><td>CUCUCGGUALCCAAGDD#GDDDAAGGCRCAAGACUGPA+APCUUGAGAD?GGGCGTPCG"CUCG<br>CCCCCGGGAGACCA</td><td>M</td><td>=</td><td>163.06333</td><td>25400.63129</td><td>25334.537</td><td>25254.5707</td><td>25005.485</td><td>24925.519</td><td>24700.444</td></tr>
<tr><td>>tdbR00000466</td><td>Val|&AC</td><td>GGUCCAAUGLUCCAGDGGDDCAAGACRPCGCCAU&ACACGGCGAAGADC?CGAGTPCG"ACCUC<br>GGUUGGAUCACCA</td><td>M</td><td>=</td><td>99.06841</td><td>24559.44837</td><td>24557.349</td><td>24477.3827</td><td>24228.297</td><td>24148.331</td><td>23923.256</td></tr>
<tr><td>>tdbR00000465</td><td>Val|CAC</td><td>GUUCCAAUALUGPAGCGGCDAUCACLPPGCCAUCACACGGCAAAG7DC?CGAGTPCG"UCCUCGG<br>UUGGAACACCA</td><td>M</td><td>=</td><td>99.06841</td><td>24477.37237</td><td>24475.273</td><td>24395.3067</td><td>24146.221</td><td>24066.255</td><td>23841.18</td></tr>
<tr><td>>tdbR00000464</td><td>Val|IAC</td><td>GGUUUCGUKGUCAAGDCGGDDAUGGCAPCUGCAUIACACGCAGAAC7DC?CCAGTPCG"UCCUG<br>GGCGAAAUCACCA</td><td>M</td><td>=</td><td>99.06841</td><td>24791.37537</td><td>24789.276</td><td>24709.3097</td><td>24460.224</td><td>24380.258</td><td>24155.183</td></tr>
</table>

**Supplemental Table 3:** Signature digestion products for detected tRNA contaminates.

| tRNA | Sequence | Mass |
|---|---|---|
| $tRNA^{PHE}$ GAA | A(Cm)U(Gm)AA(Wybutosine)A(pseudouridine)(m5C)UGp | 4164.664 |
| $tRNA^{LYS}$ 3UU | CU(mcm5s2U)UU(i6A)ACCGp | 3393.42 |
| $tRNA^{VAL}$ AAC | C(pseudouridine)U(inosine)ACACGp | 2878.371 |
| $tRNA^{TYR}$ GUA | (pseudouridine)A(t6A)A(pseudouridine)CUUGp | 2947.417 |

**Supplemental Table 4** *In silico* generated 3' digestion products lacking the terminal adenosine.

| tRNA | AC | Sequence | M | -2 | -3 |
|---|---|---|---|---|---|
| Val | IAC | AAAUCACCp | 2555.367 | 1277.684 | 843.2711 |
| Val | CAC | AACACCp | 1920.289 | 960.1445 | 633.6954 |
| Pro | NGG | CCCCCCp | 1848.255 | 924.1275 | 609.9242 |
| Leu | UAG | CUCUCACCp | 2484.317 | 1242.159 | 819.8246 |
| Leu | UAA | CAUCCUUCACCp | 3424.435 | 1712.218 | 1130.064 |
| Leu | CAA | CAACCACCp | 2530.371 | 1265.186 | 835.0224 |
| Ile | IAU | ACCACCp | 1896.278 | 948.139 | 625.7717 |
| Ala | IGC | UCCACCp | 1873.251 | 936.6255 | 618.1728 |
